# Non-invasive visualization of pH changes within the tumor-micro-environment by positron emission tomography

**DOI:** 10.1101/2024.11.21.624628

**Authors:** Jürgen Brück, Dominik Schauenburg, Seah Ling Kuan, Simeon Göttert, Benedikt Klasen, Veronika Frommberger, Kazem Ebadi Jalal, Nabil Boui, Aaron Kwiatkowski, Lisa Schake, Tobias Bopp, Tozka Bohn, Tanja Weil, Mathias Schreckenberger, Miederer Matthias

## Abstract

Slight changes in the pH value of the tumor microenvironment (TME) have crucial effects on host defense, metastatic behavior, immune regulation and cellular metabolism. Due to the high metabolic activity and insufficient perfusion of tumors, acidic metabolites often accumulate in tumors and can influence the pH of the TME. Several studies have shown that the acidity of the interstitial space and the relatively stable intracellular pH can influence the functions of cancer and stromal cells and their interaction with the extracellular matrix.

We describe radiopharmaceutical probes for positron emission tomography (PET) that exploits the increasingly used concept of pH-dependent intratumorally cleavage of PET-tracer to release functional moieties. The radiopharmaceuticals are based on [^18^F]FDG, a PET tracer, routinely used, and substituents that are pH-dependently cleaved. Utilizing preclinical models, we were able to visualize small pH differences in the acidic TME of different tumors with [^18^F]FDG-4-methoxybenzylamine ([^18^F]FDG-4MBA). *In vivo* studies were in line with in *vitro results* and showed that the hemiaminal bond between FDG and a substitute is cleaved at slightly acidic pH and leads to pH dependent radiotracer uptake. *In vivo* neutralization of the acidic extracellular tumor pH by sodium bicarbonate treatment prevents pH-dependent cleavage of [^18^F]FDG-4MBA and a resulting decrease of uptake. The determination of pH differences in acidic TME may serve as a novel marker for various questions such as regulation of the response to immunotherapies. Notably, even small pH differences in the acidic TME of different tumors, in the same *in vivo* model, could be visualized. This is the first preclinical study to show that it is possible to visualize small pH differences in the TME of different tumors in the same mouse using hemiaminal bound [^18^F]FDG. Due to the facile tracer synthesis and application, this system could be well suited for translation into clinical studies to develop new strategies of pH regulation to improve the efficacy of immunotherapy in cancer patients.

**graphical abstract:** 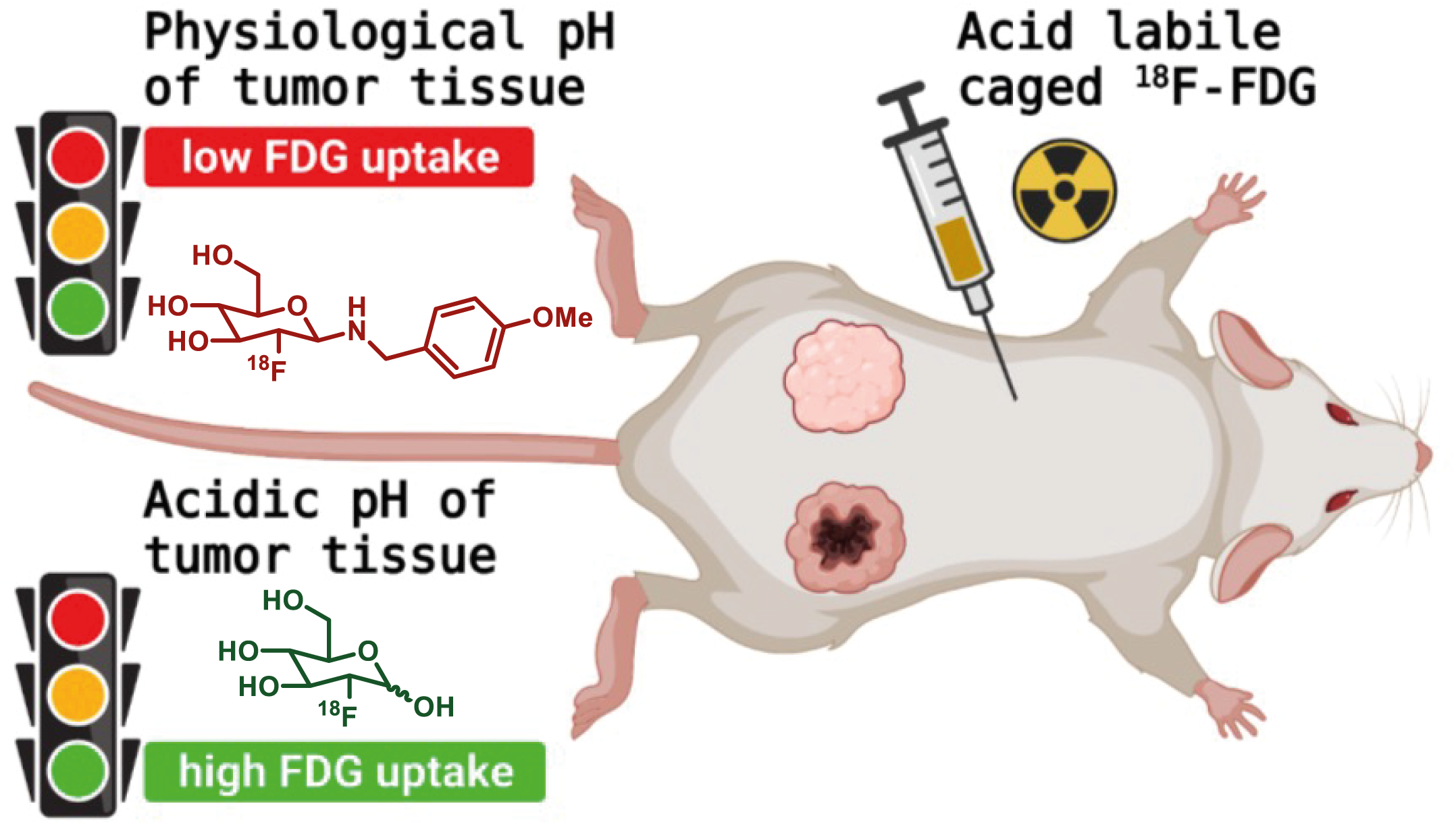

## Introduction

Minimal changes in the extracellular pH value in the tumor microenvironment (TME) have crucial effects on host defence, metastatic behaviour, immune regulation and cellular metabolism(1, 2). These phenomena of small changes of the pH in the TME, such as degranulation of the extracellular matrix, invasion of tumor cells or differentiation of tumor-associated macrophages, have been intensively investigated in various studies (3–5). Due to the high metabolic activity and insufficient perfusion of tumors, acidic metabolites often accumulate in tumors and influence the pH of the TME (6). It has been shown that the acidity of the interstitial space and the relatively stable intracellular pH can influence the functions of cancer and stromal cells and their interaction with the extracellular matrix. Changes of the intratumoral pH are spatially and temporally heterogeneous and dependent on numerous factors such as metabolism, cell density and vessel density (2, 7). Alterations in extracellular acidity have complex influences on genetic stability, epigenetics, cellular metabolism, and tumor proliferation. Different mechanisms have been proposed to explain the effects that can influence the tumor growth by a slight acidification of the TME. In this context it could be shown that the pH regulatory transport proteins NHE1 and NBCn1 can play an influence in cancer therapy depending on small pH changes in the TME. Thus, dysregulation of pH in the TME can influence and inhibit the effectiveness of cancer therapy (1, 6, 8).

The extracellular acid-base status of most normal tissue *in vivo* is relatively stable under physiological conditions and close to that of blood (9).The pH in normal tissues is usually in the range of 7.3 to 7.4, but it varies considerably under certain conditions, as in epithelium (e.g. pancreas, colon, stomach, etc.) during intense acid or base secretion and in skeletal muscle during physical activity (10–12). Small pH changes can influence the course of not only in tumor diseases. Changes in the pH value of pancreatic juice in patients with a chronic pancreatitis are associated with the inactivation of chymotrypsin C and the precipitation of lithostatin. Furthermore, numerous studies have shown that chronic metabolic acidosis is a pathognomic feature of type 2 diabetes (13, 14). Many non-tumor diseases are associated with small pH changes in injured tissue, blood or after bone fracture (10, 12, 15, 16).

Compared to the stable extracellular pH of normal tissue (pH 7.3-7.4), the pH of the TME ranging from near-neutral extracellular pH (pH 7) down to areas with intense acidification (pH 6.4) (17, 18).

This acidic microenvironment is linked to metabolic changes in tumor cells, such as the aerobic glycolysis described by Otto Warburg, leading to an increased production of acidic metabolites such as lactate and CO_2_, which are transported to the extracellular space (19, 20). Due to the reduced blood supply and higher vascular density of the tumor tissue, the metabolites accumulate in the extracellular space leads to an acidification of the TME (6, 19, 21–23).

Genetic instability and epigenetic changes are important throughout the development of most cancers. Early studies show that a strongly acidic pH can be clastogenic and induces double-stranded DNA breaks (24, 25, 26{Niu, 2022 #87, 27). The acidic TME thus provides a selection pressure that can give rise to cancer cell populations with more malignant phenotypes (19, 28, 29). Cancer cells are generally more likely to gain additional mutations, than normal cells, and this may be enhanced by extracellular acidity.

Several studies show that an acidic TME inhibits anticancer immune effector cells such as dendritic cells, T-cells, pro-inflammatory M1-like macrophages and natural killer cells (27, 30–32). At the same time, immunosuppressive components like anti-inflammatory M2-like macrophages and regulatory T-cells will be expanded and activated (33, 34). A reduction of T-cell receptor expression was demonstrated in a TME with acidic conditions (30, 35).

Furthermore, an acidic TME provides the differentiation of dendritic cells to type II DCs and thus to the production of the anti-inflammatory cytokine IL-10 and an inhibition of the pro-inflammatory cytokines IL-12 and IL-23 (31). Besides acid-induced immunosuppression, hypoxia exacerbates the negative impact of the TME on the antitumor immune response (5, 36, 37). Several preclinical studies have shown that a reduction in tumor hypoxia significantly increases T-cell infiltration in mouse prostate cancer tissue (38). Robey et al. reported that oral administration of bicarbonate increases the pH of the TME, which reduces tumor growth and metastasis in a mouse breast cancer model (39).

These published data show that tumor growth can be influenced by changing the pH in the TME and thus the antitumor effect of therapies can be improved (39, 40).

The increasingly known relationships between pH and tumor biology and the invention of new therapies addressing the TME are reasons to develop methods, which non-invasively measure pH within the tumor. Here, imaging with Positron Emission Tomography (PET) is the most attractive modality due to its reliable quantification and the lack of potential side effects of radiotracers. pH-Imaging, which can measure the pH of tumor tissue, has then the potential to guide the development and application of innovative immune modulatory therapies (36, 41). Currently used probes suffer from spatial resolution, sensitivity, and specificity, hindering accurate and comprehensive assessments (42–44). Additionally, factors such as potential interference from surrounding tissues and variability in probe distribution within the tumor can influence the reliability of pH measurements, highlighting the need for ongoing advancements in probe technologies for more precise and robust evaluations of the tumor pH environment.

We used amine nucleophiles that condense on [^18^F]FDG and form an acid-labile hemiaminal. This leads to a radiotracer that is easy to prepare using commonly available techniques and that only exhibits tumor imaging properties under acidic conditions of the TME, when hemiaminal cleavage releases [^18^F]FDG inside the tumor which then is transported and accumulated into FDG-avid tumor cells. The pK value of the hemiaminal might play a critical role for this concept and it is tailorable over a wide pKa range using appropriate amines such as aminobenzonitrile (pK 1,7), anilinium (pK 4,9), p-anisidine (pK 5,4), phenylbenzylamine (pK 9,1) and benzylamine (pK 9,2) (45). To fine-tune the pKa value and enhance nucleophilicity, electron-donating groups were introduced at the ortho- or para-position of the aromatic system (46–48). Based on available data (45), we suggest that [18F]FDG functionalized with 4-methoxybenzylamine (4-MBA) may improve imaging sensitivity and facilitate the discrimination of subtle pH variations in the tumor microenvironment (TME) across different tumor types

To evaluate the method of pH determination by hemiaminal cleavage to free [^18^F]FDG, we synthesized [^18^F]FDG-4MBA, and [^18^F]FDG-benzylamine and evaluated them in *in vitro and in vivo* models of two different murine tumor cell lines. The very aggressively growing and acidic melanoma cell line B16F10 and the colon adeno carcinoma cell line MC38, reflecting a less acidic tumor microenvironment (5, 36). We histologically characterized our B16 and MC38 tumors on day 14 after tumor cell transfer by H&E and HIF-1alpha staining. In our H&E staining we found the same proportion of neutrophils and lymphocytes in B16 and MC38 tumors. B16 tumors showed a more compact cell cluster, an onset of destruction with necrotic tumor areas and a stronger HIF-1 alpha expression than MC38 tumors (data not shown).

In contrast to [^18^F]FDG imaging, where during aerobic glycolysis [^18^F]FDG is taken up via the glucose transporter, phosphorylated to glucose 6 phosphate and accumulated in the cells, pH specific imaging with [18F]FDG-4MBA or [18F]FDG-BA depends mainly on the extracellular pH which is responsible for the cleavage of the radiopharmaceutical and only in a second step on FDG uptake by tumor cells. In this study, we were able to show that small pH differences in the TME of two different tumors in the same mouse can be visualized with an hemiaminal which is bound to [^18^F]FDG.

## Material and methods

### Mice

C57BL/6JRj mice, 6-8 weeks of age, were purchased from Janvier, All mice were housed in the animal facility (TARC) of the University Medical Center of Mainz. All animal studies were approved by the competent governmental agency Landesuntersuchungsamt Rheinland-Pfalz and were performed according to the local regulations.

### Synthesis of [^19^F]FDG-4MBA

[^19^F]FDG (5.00 mg; 27.5 μmol; 1.00 equiv.) was stirred in a solution of 4-methoxybenzylamine (10.6 mg; 82.4 μmol; 3.00 equiv.), dry DMSO (110 µL) and acetic acid (1.57 µL; 27.5 μmol, 1.00 equiv.) at 60°C for 1 hour. Subsequently, the reaction mixture was diluted with 0.5 ml of each Buffer A and Buffer B. Buffer A is an aqueous ammonium bicarbonate solution (10 mM), and Buffer B is 80% acetonitrile, 20% aqueous ammonium bicarbonate solution (10 mM) and the crude product was purified via preparative HPLC. (The gradient used for purification was as follows: 5% B for 2 minutes, increased to 30% B over 8 minutes, further increased to 40% B in 3 minutes, and finally reached 100% B over 4 minutes). The product was obtained as a colorless powder after lyophilization: 5.47 mg (18.2 μmol; 66 %).

### HPLC Purification of [^19^F]FDG-4MBA

The reaction mixture was purified on a reverse phase high-performance liquid chromatography. The column used was an Agilent Eclipse XDB-C18, 5 um, 9.4 x 250 mm from Agilent Technologies (USA, Santa Clara). Buffer A: aqueous ammonium bicarbonate solution (10 mM), Buffer B: 80% acetonitrile, 20% aqueous ammonium bicarbonate solution (10 mM), at a flow rate of 1 mL/min.

### Nuclear magnetic resonance (NMR) spectroscopy

The release of [^19^F]FDG from [^19^F]FDG-4MBA was monitored using ^1^H nuclear magnetic resonance (NMR) spectroscopy (Bruker Avance 700 NMR spectrometer). Chemical shifts are expressed in parts per million (ppm) and are referenced to residual non-deuterated solvent signals (DMSO). 1 mg [^19^F]FDG-4MBA was dissolved in 500 µL of deuterated phosphate buffer (pH 7.4, 6.9 or 6.5), and 1 mg DMSO was added as a reference. Over a period of 20 hours, a ^1^H spectrum was recorded every 15 minutes.

### Synthesis of [^18^F]FDG-4MBA and [^18^F]FDG-BA

[^18^F]FDG was obtained from Advanced Accelaration Application Germany. Synthesis of [^18^F]FDG-4MBA and [^18^F]FDG-BA were carried out on a SynChrom R&D module from Elysia Raytest. First, [^18^F]FDG was azeotropically dried using acetonitrile under vacuum and a stream of nitrogen. Afterwards, a solution of 0.5 M 4-methoxy-benzylamine or benzylamine and 0.25 M acetic acid was added to the dried [^18^F]FDG. The reaction mixtures were incubated for 30 minutes at 80°C under stirring. The reaction mixture was diluted, and the crude product subsequently purified via preparative HPLC (isocratic; 15% acetonitrile). The collected fractions were diluted with water and passed through a C18 Sep Pak Plus cartridge (Waters GmbH, Germany). Finally, the trapped products [^18^F]FDG-4MBA and [^18^F]FDG-BA were eluted with ethanol and formulated with bicarbonate buffer.

### PET

The injections of [^18^F]FDG-4MBA, [^18^F]FDG-BA or [^18^F]FDG were performed on anesthetized animals with 2.0% isoflurane (Abbott, Wiesbaden, Germany) evaporated in oxygen at a flow rate of 0.5 L/min. Immediately before the injection of [^18^F]FDG, [^18^F]FDG-4MBA or [^18^F]FDG-BA body weights of the mice were measured. In total, ∼5 MBq PET-tracer dissolved in 100 µL 0.9% NaCl solution was injected i.v. Whole body PET scans were performed after injection of [^18^F]FDG-4MBA, [^18^F]FDG-BA or [^18^F]FDG after 45 minutes using a dedicated Focus 120 small-animal PET scanner (Concorde Microsystems/ Siemens Preclinical Solutions, Knoxville, USA), for 12 min while the mice were kept under anaesthesia. A reconstruction was performed in microPET Acquisition Workplace V02.3200 with OSEM3D followed by map. The reconstructed voxel size was 0.432×0.432×0.796 mm^3^. The percentage of injected dose per cubic centimetre (%ID/cm³) values were calculated as follows: mean activity in VOI/(injected activity·10^6^)/100.

### Biodistribution

In addition to *in vivo* PET data, organs were analysed by γ-counting *ex vivo* in separate groups, 55 min after tracer injection. A standard solution with a known radioactivity served as reference to calculate the percentage of injected dose per gram of tissue (%ID/g). The tubes containing the standard solution, and the organs were measured via Wizard2 automated γ-counter (Perkin Elmer) at an energy window of 350 to 650 keV. The resulting decay corrected counts per minute were then first normalized to the injected dose with the help of the standard solution and to the respective weight of the organ to obtain %ID/g.

### Cell lines and subcutaneous tumor induction

B16F10 and MC38 cells were maintained in Dulbecco’s modified Eagle’s medium (ThermoFisher Scientific, Germany) containing 10% heat-inactivated fetal bovine serum (FBS), 2 mM L-glutamine, 1% penicillin and streptomycin (all from ThermoFisher Scientific, Germany). Tumor cells were taken from the log phase of *in vitro* growth (70% confluency) and transferred into the left (B16; 2×10^5^ cells/mouse) and right (MC38: 1×10^6^ cells/mouse)) flank of the mice. Tumors were visible after 10 days and grew in comparable size in both flanks of the experimental animals. The size of the inoculated tumors was monitored daily for the duration of the experiments

### Sodium Bicarbonate Treatment

Mice were supplemented with 200 mM sodium bicarbonate in their drinking water, control mice received normal drinking water. Bicarbonate feeding began 10 days before tumor inoculation. PET-imaging was performed on days 10-14 after tumor cell transfer.

### Cellular tracer uptake study

For cell uptake experiments, B16 or MC38 cells were seeded at a density of 4 × 10^6^ cells in Falcon tubes containing PBS including 5%FCS with a different pH value ranging from 6.5 to 7.4. Then 0.1 MBq of [^18^F]FDG-4MBA or [^18^F]FDG was added to each sample. Cells were incubated at 37 °C, 5% CO_2_ for 20 min. Following incubation, the cells were washed three times with PBS. The tubes containing the different samples were measured via Wizard2 automated γ-counter (Perkin Elmer) at an energy window of 350 to 650 keV to quantify the amount of ingested tracer.

### HPLC Degradation Assay

After purification by semipreparative HPLC and subsequent trapping on a C18 cartridge (Waters, Sep-Pak^TM^ Plus) the products [^18^F]FDG-4MBA and [^18^F]FDG-BA were eluted using ethanol and formulated either with sodium carbonate buffer pH 8 or sodium phosphate buffer pH 6.5. To analyse the cleavage of [^18^F]FDG-4MBA and [^18^F]FDG-BA at different pH, the product solutions were incubated and measured for three to four times in an analytical HPLC system at intervals of 30 minutes

### Ex vivo pH measurement in tumors with unisense microelectrode system

pH-microelectrodes for *ex vivo* pH measurement of tumors was purchased from Unisense A/S (Denmark). The signals were acquired by a microsensor multimeter (Unisense, Denmark) interfaced with a computer to record data. Microelectrodes were calibrated with three standard solutions. To offset the disturbance of the electric field, calibrations of pH and ORP microelectrodes were performed under the same potential as that of the working electrode. Tumor-bearing animals were euthanized on day 14 after tumor inoculation, B16 and MC38 tumors were dissected, and the pH was determined directly with the unisense microelectrode system.

### Statistics

All data were analysed and plotted using GraphPad Prism 8 software. Statistical analyses were performed by using the Mann-Whitney test, the Wilcoxon test or the Tukey’s post-hoc test after One-way ANOVA as indicated. Values of P<0.05 were considered significant.

## Results

### Synthesis of [^19^F]FDG-4MBA

Hemiaminal formation using 4-MBA and [^19^F]FDG under acidic conditions (formic acid) results in [^19^F]FDG-4MBA, which could be isolated in 66% yield by preparative HPLC.

### Release of condenses [^19^F]FDG monitored by ^1^H nuclear magnetic resonance (NMR)

Release of [^19^F]FDG from [^19^F]FDG-4MBA was monitored by ^1^H nuclear magnetic resonance (NMR) spectroscopy under aqueous buffered conditions. Aromatic protons of the 1,4-substituted ring undergo a significant downfield shift after hemiaminal hydrolysis, and the time courses of release were recorded at different pH values by NMR. Under physiological conditions (pH 7.4) the decaying time, at which 50 % is released, is 3 hours (Fig. 1A). Mimicking the pH conditions of a slightly acidic TME (pH 6.9), a shortening to only about 1 hour was found (Fig. 1B). This observation fits well to the radioactive decay time of the F-18 analogue ([^18^F]FDG-4MBA). According to the half-life of hot [^18^F]FDG (110 min), this means that at pH 7.4 two thirds of [^18^F]FDG are still present as a condenses [^18^F]FDG complex, whereas at pH 6.9 only one third remains. This is even more pronounced at pH 6.5, where 50% is released after 45 minutes and almost 80 % after 110 min (Fig. 1C). Figure 1D shows the stability of the hemiaminals as a function of the pH value. We found that at pH 7.4 there is twice as much bound [^19^F]FDG as at pH 6.5 after 1 hour. After 2 hours, this trend becomes even more significant: At pH 7.4 there is about three times more [^19^F]FDG hemiaminal compared to pH 6.5.

**Figure 1:**
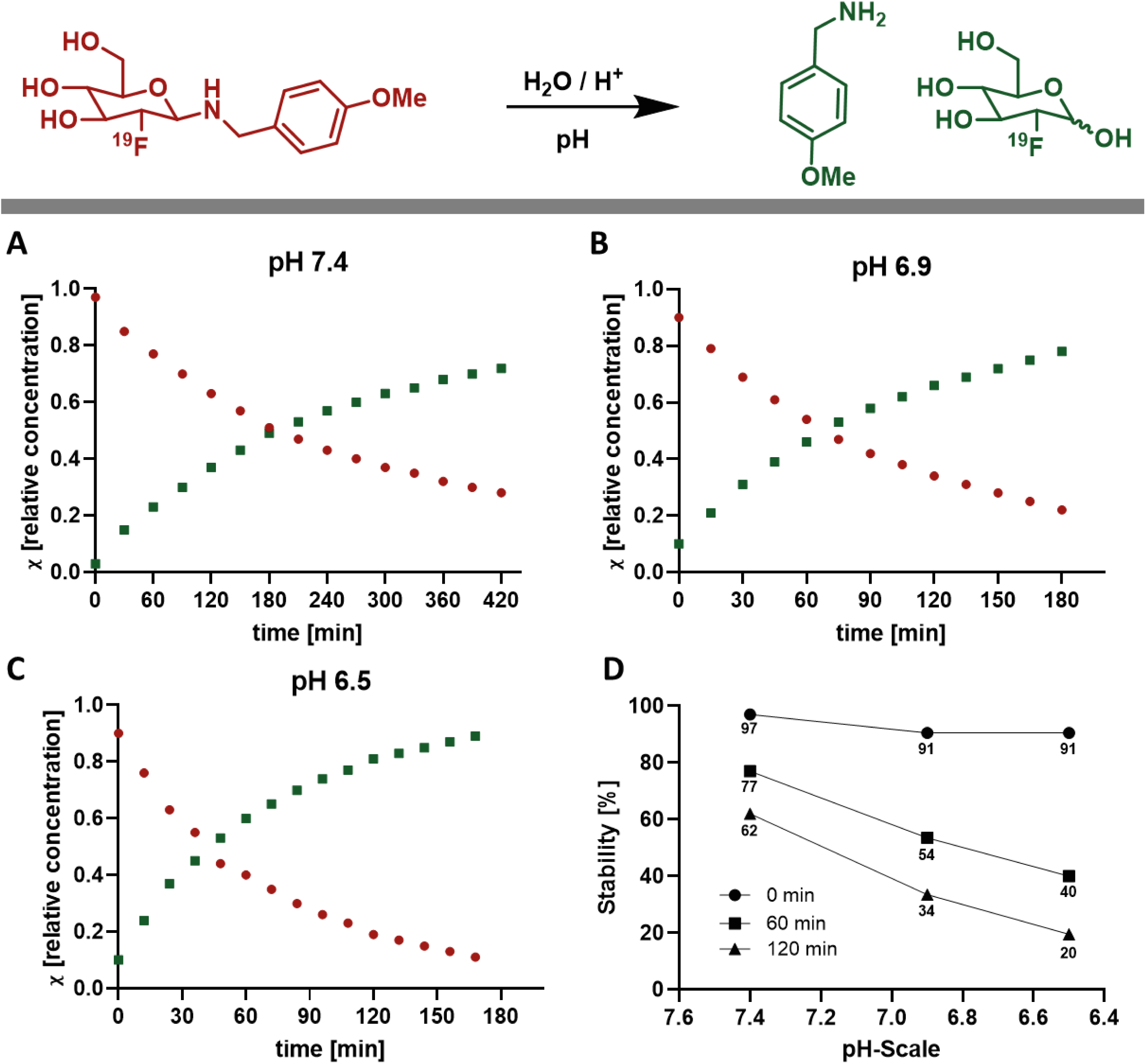
Synthesis of [^19^F]FDG-4MBA. *Time dependent release studies of cold* [^19^F]FDG-4MBA *monitored by 1H nuclear magnetic resonance (NMR) spectroscopy at different pH values*.

### Synthesis of [^18^F]FDG-4MBA, [^18^F]FDG-BA and [^18^F]FDG-4PBA

The synthesis of [^18^F]FDG-4MBA and [^18^F]FDG-BA were carried out in one-step Elysia-Raytest synchron module by incubation of [^18^F]FDG with 4-methoxy-benzylamine or benzylamine (Fig. 1A), at 80°C for 30 min. The synthesized hemiaminals were purified using a preparative HPLC system. Preparation and purification of [^18^F]FDG-4-phenylbenzylamine was not pursued further in this study due to the insufficient solubility of 4-phenylbenzylamine and the low product yields. In contrast, sufficient [^18^F]FDG-4MBA and [^18^F]FDG-BA could be eluted at 6-8 minutes from the C18 10µm column whereas unconjugated free [^18^F]FDG elutes at 2-4 minutes (Fig 2A, B). The average radiochemical conversion, based on integration of HPLC peaks, amounted to 69 ± 7% at [^18^F]FDG-4MBA and 64 ± 4% [^18^F]FDG-BA.

**Figure 2:**
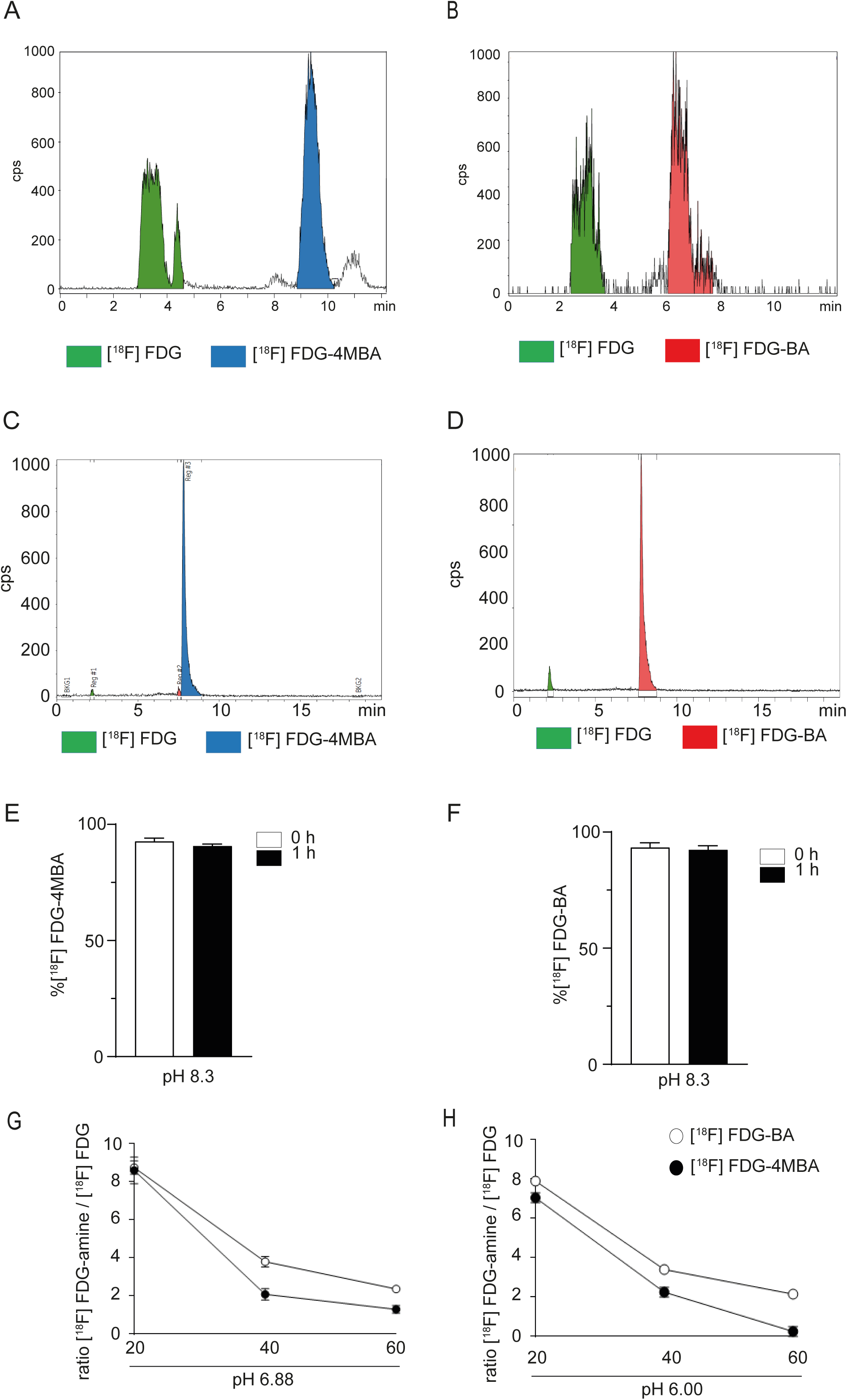
Synthesis of [^18^F]FDG-4MBA and [^18^F]FDG-BA. Synthesis of [^18^F]FDG-4MBA and [^18^F]FDG-BA were carried out in one-step Elysia-Raytest synchron module by incubation of [^18^F]FDG with 4-methoxy-benzylamine (**A**) or benzylamine (**B**). The synthesized hemiaminals were purified using a preparative HPLC system. [^18^F]FDG-4MBA and [^18^F]FDG-BA could be eluted at 6-8 minutes from the C18 10µm column whereas unconjugated free [^18^F]FDG elutes at 2-4 minutes. Data show representative syntheses of [^18^F]FDG-4MBA and [^18^F]FDG-BA. (**C** and **D**) The purity of [^18^F]FDG-4MBA and [^18^F]FDG-BA was determined using an analytical HPLC system. The chromatogram showed high purity (< 95%) of both products (elution at 8 minutes) and no free [^18^F]FDG. Data show representative HPLC chromatograms. To evaluate the stability [^18^F]FDG-4MBA and [^18^F]FDG-BA were eluted in buffers with increasing acidity pH 8 and pH 6.88 and injected at intervals of 20 minutes into the analytical HPLC system. Both radiopharmaceuticals eluted in a buffer with a pH of 8.3 showed no hydrolysis over a time of one hour (**E** and **F**). Bars represent mean ± SEM of four independent experiments. (**G** and **H**) [^18^F]FDG-4MBA and [^18^F]FDG-BA were eluted into a buffer with pH 6.88 or pH 6.00. Hydrolysis of both radiopharmaceutical is presented as a ration between free [^18^F]FDG and [^18^F]FDG-4MBA or [^18^F]FDG-BA. Data represent mean ± SEM of three independent experiments.

To confirm the purity of [^18^F]FDG-4MBA and [^18^F]FDG-BA for following experiments, both products were injected into an analytical HPLC system. The chromatogram showed high purity (< 95%) of both products (elution at 8 minutes) and no free [^18^F]FDG (Fig. 2 C, D). To further evaluate the stability of [^18^F]FDG-4MBA and [^18^F]FDG-BA at different pH values *in vitro*, we investigated their release times *versus* the pH of the applied buffer. [^18^F]FDG-4MBA and [^18^F]FDG-BA respectively were eluted in buffers with increasing acidity and were injected at intervals of 20 minutes into the analytical HPLC system. [^18^F]FDG-4MBA and [^18^F]FDG-BA eluted in a sodium-bicarbonate-buffer (pH 8.3) showed no hemiaminal hydrolysis over more than one hour (Fig. 2E, F). To simulate the hydrolysis of our [^18^F]FDG-conjugates under a pH value corresponding to the TME *in vivo*, we eluted [^18^F]FDG-4MBA and [^18^F]FDG-BA into a buffer with a pH of 6.88. Fig. 2G shows the ratio between the [^18^F]FDG - conjugates and free [^18^F]FDG over time. In contrast to [^18^F]FDG-BA, [^18^F]FDG-4MBA shows a stronger hydrolysis at a pH of 6.88 like the pH of the TME *in vivo* over time. Under an even more acidic buffer with a pH of 6.0, the difference in the hydrolysis of [^18^F]FDG-4MBA and [^18^F]FDG-BA is even more marked (Fig. 2H).

Our data shows that in a pH range between 6 and 7, an hemiaminal consisting of [^18^F]FDG and 4MBA is easily hydrolysed and therefore is well suited to discriminate pH differences in the TME *in vivo* (Fig. 2G).

### Tumor cell uptake of [^18^F] FDG-4MBA and [^18^F] FDG-BA under acidic culture conditions in vitro

To simulate pH differences in the TME and their effect on the pH-dependent [^18^F]FDG release from [^18^F]FDG-4MBA and [^18^F]FDG-BA *in vitro*, tumor cells (B16 and MC38) were cultured under increasing acidity of the cell culture medium from 7.6 to 6.4. Tumor cells were processed with the same level of activity of [^18^F]FDG-4MBA or [^18^F]FDG-BA and as a control for a pH independent cellular enrichment [^18^F]FDG was used. Both cell lines showed a pH dependent enrichment of [^18^F]FDG-4MBA and [^18^F]FDG-BA (Fig. 3A-D), whereas unmodified [^18^F]FDG did not show a pH dependent uptake (Fig. 3E, F).

**Figure 3:**
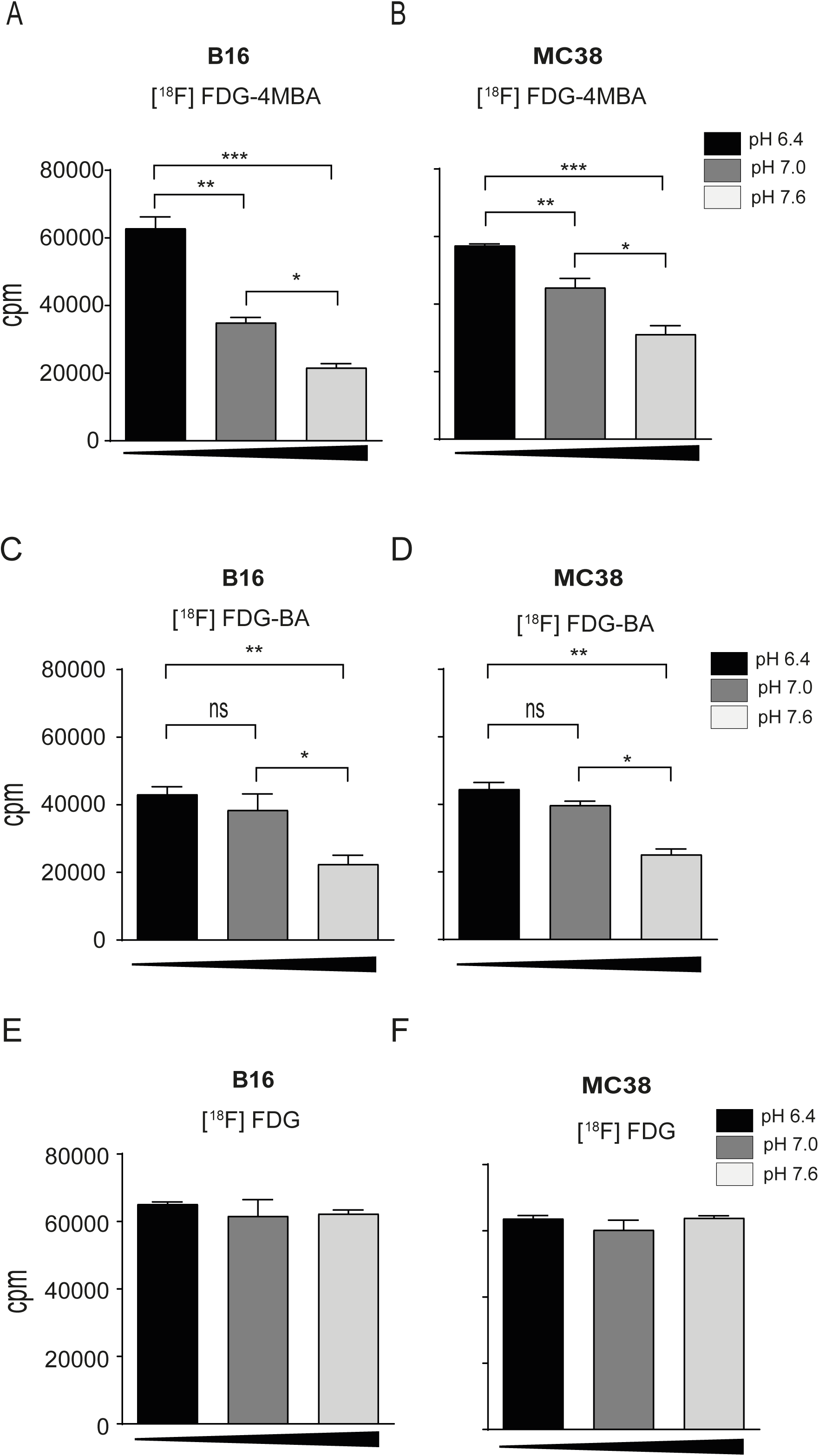
Cellular enrichment of [^18^F]FDG-4MBA and [^18^F]FDG-BA uptake under acidic culture conditions in vitro. (**A**-**F**) tumor cells (B16 and MC38) were cultured under increasing acidity of the cell culture medium from 7.6 to 6.4. Tumor cells were processed with the same level of activity of [^18^F]FDG-4MBA (**A**-**B**) or [^18^F]FDG-BA (**C**-**D**) and as a control for a pH independent cellular enrichment [^18^F]FDG (**E**-**F**) was used. The cellular uptake of [^18^F]FDG-4MBA and [^18^F]FDG-BA were measured via Wizard2 automated γ-counter (Perkin Elmer). Bars represent the mean ± SEM of four independent experiments.

In our cellular *in vitro* assay to simulate the pH range of the TME *in vivo*, we found no measurable difference in the cellular uptake of [^18^F]FDG-BA in a pH range corresponding to the TME *in vivo* of 6.4-7.0 (Fig. 3C, D). These data show that [^18^F]FDG-4MBA is similar to the already described [^18^F]FDG-BA (45) suited to determine pH changes in a range of 7.0 to 7.6 (Fig. 3C, D) and better suited in pH range of the TME between 6.4 to 7.0.

### Determination of [^18^F] FDG-4MBA distribution in naïve mice compared to [^18^F]FDG

To confirm the relevance of our *in vitro* findings *in vivo*, we injected [^18^F]FDG-4MBA and [^18^F]FDG in C57BL/6 wildtype mice and quantified the uptake by PET imaging and *ex vivo* biodistribution analysis. The highest accumulation of [^18^F]FDG was observed in the heart (14,8±2,0 % ID/cm^3^). In mice after injection of [^18^F]FDG-4MBA, we determined a significantly lower uptake in the heart (5.0±1.2% ID/cm^3^) indicating that unspecific background accumulation either from intact initial tracer or from unspecifically released FDG is present (Fig. 4A-B). However, a reduction to one third can well be considered strong enough to shift the focus of the molecular basis of the imaging strongly from FDG uptake to pH measurement. In addition, we confirmed the non-invasive PET imaging data by *ex vivo* biodistribution (Fig.4C).

**Figure 4:**
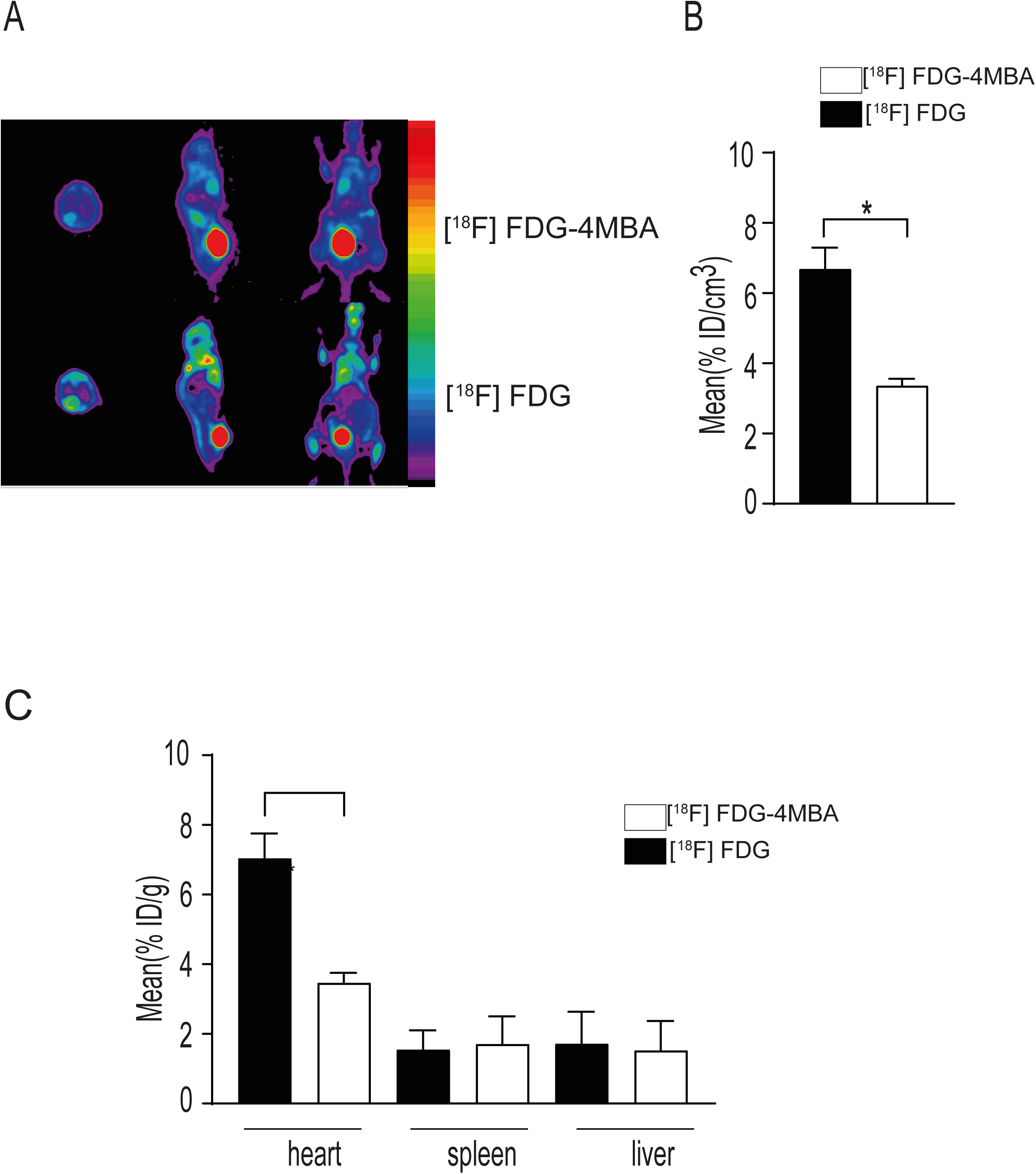
Determination of [^18^F]FDG-4MBA distribution in naïve mice compared to [^18^F]FDG. [^18^F]FDG-4MBA and uptake [^18^F]FDG was quantified by PET imaging and biodistribution. Images from single mice are depicted in (A). Pooled [^18^F]FDG, [^18^F]FDG-4MBA PET (B) and biodistribution (C) data from three independent experiments are shown (n=5). Box plots show the median with 25th and 75th interquartile range (IQR) and whiskers indicate 1.5 x IQR (*P<0.05). Data in were analysed using Tukey’s post hoc test after One-way ANOVA.

### Determination of the TME pH range in vivo

To test the possibility of quantifying small differences in the pH range of a TME with PET, we used a mouse model with two different tumors, each of them has a TME with a different pH range. As shown in previous studies, a B16 mouse-melanoma shows a more acidic TME than the MC38 adenocarcinoma. To be able to estimate the pH differences of the TME in B16 and MC38 tumors, we determined the pH value in both TMEs on day 14 after tumor cell transfer, using a unisense pH electrode system. As described in numerous studies, an MC38 adenocarcinoma has a less acidic TME (pH 6.7) than a B16 (pH 6.3) melanoma of the same size (Fig. 5A). To demonstrate that we are also able to measure pH changes in the TME as a basis for subsequent imaging with a pH dependent cleavable radiopharmaceutical, we fed our experimental mice with bicarbonate prior tumor cell transfer. Bicarbonate feeding is a classical protocol to influence the pH in the TME (3). In contrast to mice from the control group, whichonly received drinking water (Fig. 5A), bicarbonate feeding resulted in a more alkaline TME in both B16 melanomas (pH 7.2) and MC38 adenovcarcinomas (pH 7.5) (Fig. 5B).

**Figure 5:**
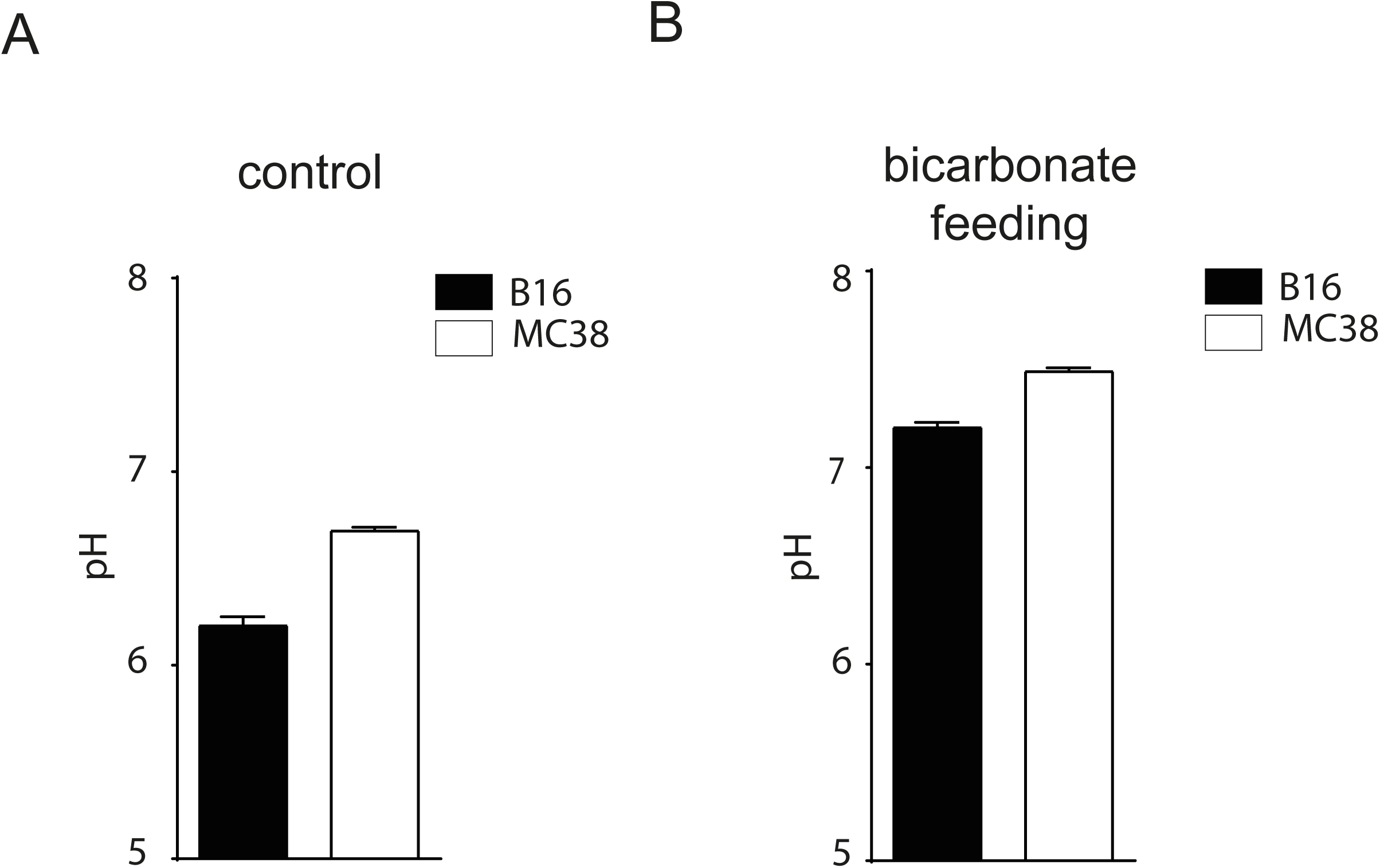
Determination of the TME pH range *in vivo*. (A) The pH value in the TME of the B16 melanoma (left mouse flank) and the MC38 adenocarcinoma (right flank) on day 14 after tumor cell transfer was determined using a unisense pH electrode system (A). (B) Mice received bicarbonate or control water on day 14 after tumor cell transfer, the pH values in the different TMEs were determined using a unisense pH electrode system. Data from two independent experiments are shown (n=8).

### Non-invasive discrimination of tumors with different pH in the TME by PET

To investigate whether it is possible to discriminate small differences in acidity in the TME we used the two tumor models (B16 and MC38) with different acidic TME in the same mouse. First, we injected [^18^F]FDG as a marker of glucose metabolism in in B16- and MC38 bearing mice. On day 12 after tumor cell inoculation, no differences in [^18^F]FDG uptake between B16 (6.61±1.6% ID/cm^3^ and MC38 (7,20±1.40% ID/cm^3^) were determined in tumors with the similar volumes (B16: 604 mm^3^, MC38: 652 mm^3^; Fig. 6A,B). *Ex vivo* biodistribution analysis confirmed the non-invasive *in vivo* [^18^F]FDG-PET imaging results, as we determined the same [^18^F]FDG uptake in B16 (7.4±1.7% ID/g) and MC38 (8±1.4% ID/g) tumors, from the same mice (Fig. 6C). Therefore, in line with the *in vitro* results no major differences between the tumors regarding FDG uptake are present. Next, we evaluated the radiopharmaceuticals [^18^F]FDG-4MBA and [^18^F]FDG-BA in these tumor bearing mice for their ability to discriminate the different TMEs in a direct comparison of both tumors. On day 12 after tumor cell inoculation we determined a significantly higher [^18^F]FDG-4MBA uptake in B16 tumors (5.6±0.5% ID/cm^3^) compared to MC38 tumors (4.0±0.4% ID/ cm^3^) (Fig. 6D, E). Again, *ex vivo* biodistribution analysis confirmed [^18^F]FDG-4MBA PET imaging results, as we determined a significantly higher [^18^F]FDG-4MBA uptake in B16 tumors (5.8±0.7% ID/g) compared to MC38 tumors (3,9±0.5% ID/g) (Fig. 6F). In contrast, tumors displayed also uptake of [^18^F]FDG-BA, but the differences in acidity levels in the TME of tumors could not be quantified with [^18^F]FDG-BA. Here we determined differences in the [^18^F]FDG-BA uptake between B16 (4.3±0.4% ID/cm^3^) and MC38 (4.4±0.6% ID/cm^3^) tumors (Fig. 6G,H). *Ex vivo* biodistribution of tumor-bearing mice after application of [^18^F]FDG-BA confirms our non-invasive imaging data of B16 (4.4±0.35% ID/g) and MC38 (4.3±0.2% ID/g) (Fig. 6 I). To better elucidate the differences in the quantification of small pH differences in acidic TME with [^18^F]FDG-4MBA and [^18^F]FDG-BA, we normalized the uptake of both radiopharmaceuticals to the enrichment of normal [^18^F]FDG in B16 and MC38 tumors (Fig. 6B, E, H). When comparing the acidic TME of B16 tumors with the less acidic TME of MC38 tumors, there is a difference of 29% between B16 (84%) and MC38 (55%) based on the normalized tracer uptake with [^18^F]FDG-4MBA. In contrast, if we compare the normalized uptake of [^18^F]FDG-BA in PET imaging of the TME between B16 (65%) and MC38 (61%), only a difference of 4% is quantifiable. These data show that the nucleophilic substituent bound to [^18^F]FDG influences the sensitivity of the radiopharmaceutical for non-invasive pH imaging.

**Figure 6:**
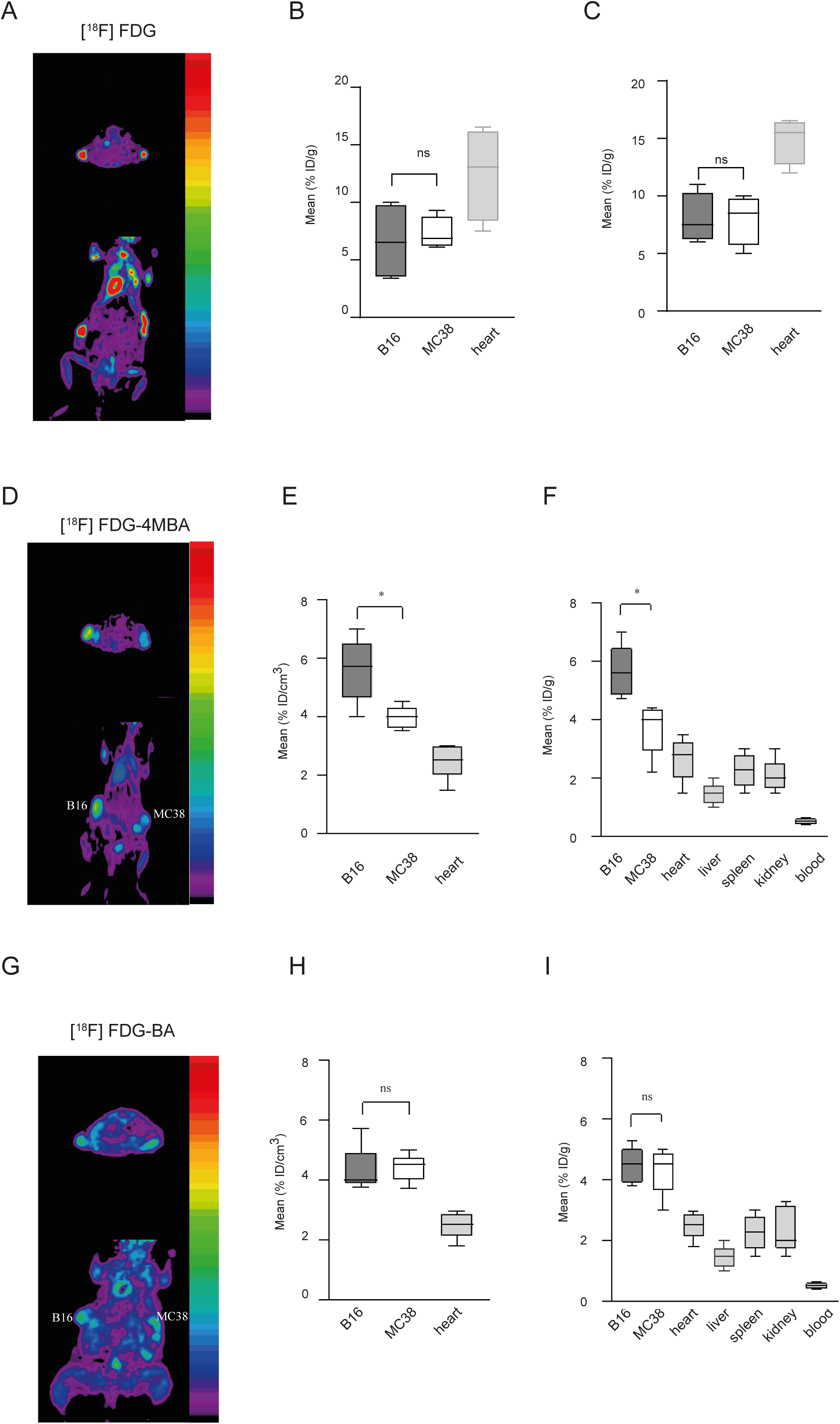
Non-invasive discrimination of tumors with different pH in the TME by PET. Mice received B16 and MC38 tumor cells and imaged on days 12 after tumor cell transfer. [^18^F]FDG, [^18^F]FDG-4MBA and [^18^F]FDG-BA uptake in the TME of B16 and MC38 tumors were quantified by PET imaging and biodistribution. Images from single mice are depicted in (A, D, G)Pooled [^18^F]FDG (B), [^18^F]FDG-4MBA (E), [^18^F]FDG-BA (F) and biodistribution (C, F,I) data from five independent experiments are shown ( n = 20).Box plots show the median with 25th and 75th interquartile range (IQR) and whiskers indicate 1.5 x IQR (*P<0.05). Data in were analysed using Tukey’s post hoc test after One-way ANOVA.

In order to be able to show the different uptake of [^18^F]FDG-4MBA and [^18^F]FDG-BA at different pH values in more detail, we normalized the uptake of both radiopharmaceuticals to the liver uptake. Numerous published studies relate tracer uptake in different tissues to tracer uptake in the liver. In our experiments, the liver showed the lowest tracer accumulation with an equal accumulation of [^18^F]FDG-4MBA and [^18^F]FDG-BA (Fig. 6F, I). [^18^F]FDG-4MBA and [^18^F]FDG-BA showed an uptake of (1.5±0.2% ID/g) and (1.7±0.3% ID/g) in the liver. When comparing the acidic of the different TMEs. B16 tumors showed a 4-fold higher uptake of [^18^F]FDG-4MBA compared to the uptake in MC38 (2.5-fold) tumors, related to the uptake in the liver, which was set to 1 after normalization.

With these data normalized to the liver uptake and the uptake of [^18^F]FDG, it can be shown that the functionality of these form of pH cleavable pet tracer consisting of [^18^F]FDG and a cleavable hemiaminal seems to depend on the properties of the conjugated amine.

### Alkalinisation of the TME prevents the quantification of small differences of the acidity in TME

We examined the possibility of visualizing small differences in the acidic TME with [^18^F] FDG-4MBA after alkalisation by bicarbonate treatment. Treatment with sodium bicarbonate has been used in several studies as a possible tumor therapy through alkalization of the TME by an average of 0.3–0.8 pH units, without significant effects on systemic pH. (45). Mice were treated with bicarbonate in drinking water for 10 days prior tumor transfer. On day 12 after tumor inoculation, we determined a lower [^18^F]FDG-4MBA uptake in B16 tumors from sodium bicarbonate treated mice (2.7±0.7% ID/cm^3^) when compared to B16 tumors from non-sodium bicarbonate treated control mice (5.1±0.6% ID/cm^3^) (Fig. 7A-C; 6D-F). *Ex vivo* biodistribution analysis of the experimental mice confirmed our non-invasive *in vivo* [^18^F]FDG-4MBA PET imaging results, as we determined a lower [^18^F]FDG-4MBA uptake in sodium bicarbonate-treated mice (2.2±0.5% ID/g) compared to non-treated littermates (4.95±0.7% ID/g) Fig. 7C;6F).

**Figure 7:**
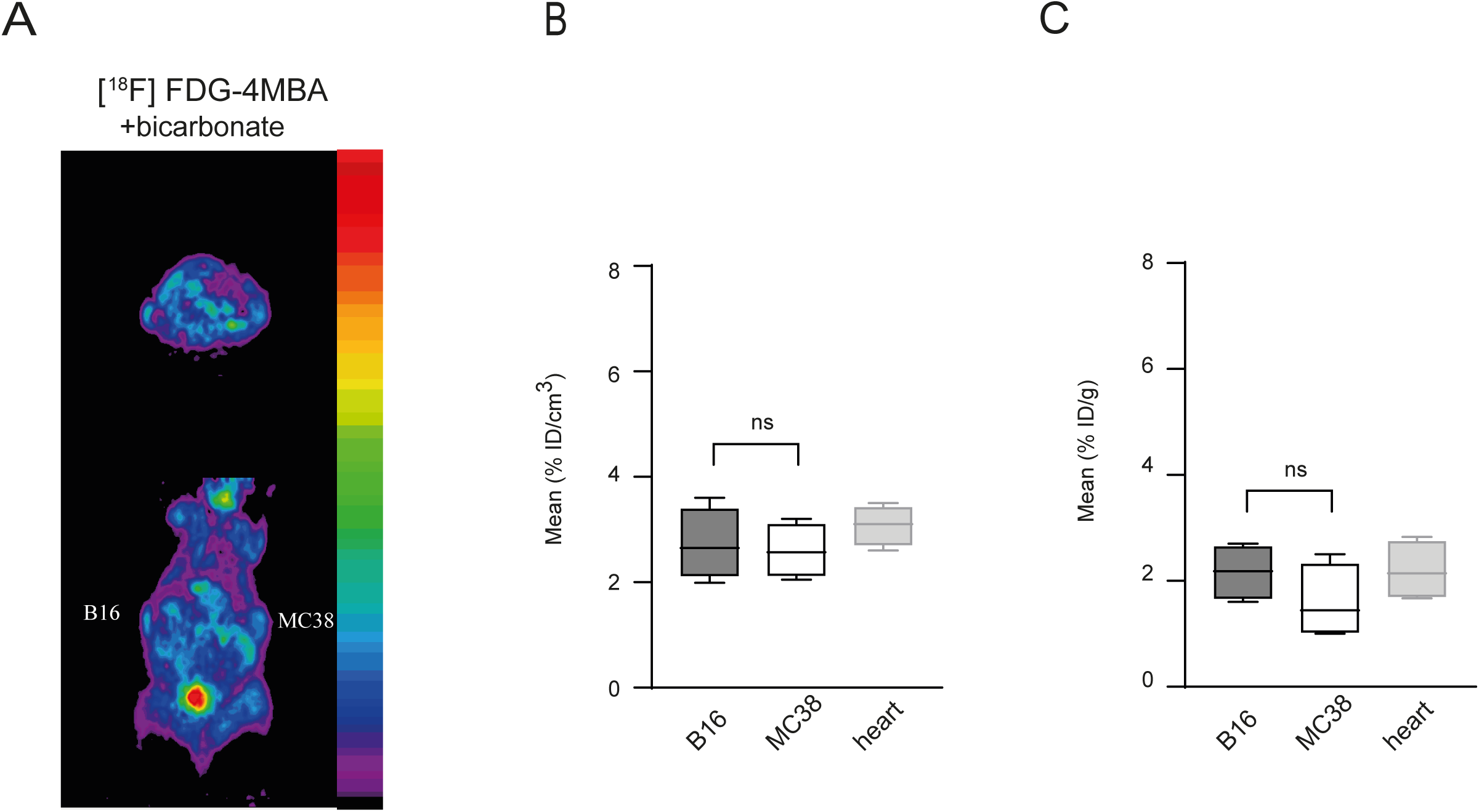
Alkalinisation of the TME prevents the quantification of small differences of the acidity in TME with [^18^F]FDG-4MBA. Mice received bicarbonate or control water and received B16 and MC38 tumor-cells. Mice were imaged on day 12 after cell transfer. [^18^F]FDG-4MBA uptake in TME of B16 and MC38 tumors was quantified by PET imaging and biodistribution. Images from single mice are depicted in (A). Pooled [^18^F]FDG-4MBA PET (B) and biodistribution (C) data from three independent experiments are shown (n=9). Box plots show the median with 25th and 75th interquartile range (IQR) and whiskers indicate 1.5 x IQR (*P<0.05). Data in (A to E) were analysed using Tukey’s post hoc test after One-way ANOVA.

## Discussion

Several different non-invasive methods for determining the pH in the TME have been described. In preclinical experiments, fluorescence probes for imaging the pH value in tissue have been developed over the past years. These techniques are very easy to use due to their high selectivity, sensitivity and fast response with relatively simple instrumentation (49) Nevertheless, these pH probes often have several limitations, such as interference from instrument efficiency, high fluorescence background due to autofluorescence of haemoglobin, artefacts in photoconversion and complicated synthesis. Thus, these probes and imaging setups are less suitable for routine clinical applications due to their low penetration depth into the tissue and inefficient quantification. Magnetic resonance imaging (MRI) is used both in clinical and basic biomedical research. However, MRI detection of pH changes poses a technical challenge. One problem with pH measurements using H-MRI sequences is that there is no endogenous pH indicator for this method. Therefore, exogenous substances that react sensitively to pH changes with chemical shifts must be administered for a pH measurement with H-MRI sequences (50).

Different radiopharmaceutical based approaches have already been used to measure the pH value. When using e.g. [^11^C]-labelled dimethyloxazolidinedione (DMO) for pH measurement, the function is based on the distribution of the neutral and ionized forms. It is assumed that the blood-brain barrier for the neutral form of the probe (HMDO) can pass through it. In the brain, pH-dependent DMO is formed from HDMO, which can no longer pass the blood-brain barrier. The method is based on several assumptions, such as the homogeneous distribution of the tracer in the tissues. In addition, the DMO distribution depends on the pH gradient of the plasma membrane and the fractional volume of the intra- and extracellular space, which further complicates the analysis and restricts it to application within the brain. Both [^11^C]O_2_ and DMO measure relative rather than absolute pH, and the short half-life of [^11^C] (∼20 min) limits its use as an on-site cyclotron is required (51, 52). Flavell et. al. was able to show that a nucleophilic acid-labile substitution of an amine to [^18^F]FDG can image tumors based on small pH differences (45). These [^18^F]FDG hemiaminals function via a two-step mechanism in which the acid-labile [^18^F]FDG-[nucleophilic substituent] decompose to form the common radiotracer [^18^F]FDG and again the nucleophilic substituent, which is subsequently accumulated by glucose-affinity cells. Thus we were able to visualize tumors with an acidic TME, as already described by Flavell et al. In our study, we were able to show that [^18^F]FDG-4MBA can be used to discriminate small differences in pH of tumors that might be essential for pH directed tumor therapy.

Due to the poor solubility, the difficult synthesis of [^18^F]FDG-4PBA and the poor yield of these radiopharmaceutical, we decided to use 4-methoxybenzylamine as an amine, which has sterically good properties for the visualization of small pH differences of the TME. In contrast to 4PBA, 4MBA is soluble in water and our data obtained in this study show that it is very well suited to visualize small pH differences both *in vitro* and *in vivo*. Benzylamine is also very soluble in water and the synthesis also provides high yields of [^18^F]FDG-BA. With the slightly larger substitute the resulting radiotracer [^18^F]FDG-4MBA showed similarities with [^18^F]FDG-BA. However, small pH differences *in vitro* and *in vivo* can be visualized better with 4MBA conjugated to [^18^F]FDG than with [^18^F]FDG-BA.

Flavell et al. also hypothesize that the hydrolysis rate of [^18^F]FDG-amines depends on the utilized amines, whose pKa should correspond to the pH of the TME.

The pH-dependent hydrolysis of glycosylamines was also described by Isbell S.H. and Frush H. L. In these studies, the authors were also able to show that these effects are only possible in a limited pH range(53, 54). An alternative option to glycosylamines are small polymers that are degraded under acidic conditions. These are used in new therapeutic approaches to trigger the release of drugs as soon as the drug carrier has reached the slightly acidic tumour environment or after the drug carrier has been taken up by the cells, which leads to the localization of the polymer in the acidic endosomes and lysosomes (55). In the initial stage of heminaminal hydrolysis, reversible protonation occurs, wherein the pKa value of the nitrogen plays a crucial role. Ideally, this pKa value should align with the slightly acidic pH of the TME to facilitate protonation, allowing for subsequent cleavage of the radiopharmaceutical into free [^18^F]FDG and the amine. This pH-dependent process suggests a potential mechanism for selective activation within the acidic TME compared to physiological pH conditions. Immunotherapies and therapies with checkpoint inhibitors have revolutionized the treatment of cancer (56–58). The effectiveness of these therapies is often limited by the hypoxic and acidic TME, which can influence the expression of checkpoint molecules. This leads to T-cell exhaustion and enables the tumor immune system to be bypassed (36). Numerous data have shown that by determining the acidity of the TME, predictions can be made about the aggressiveness of tumor growth, metastatic behavior and the migration and differentiation of immune cells (6, 8, 18). By determining the acidity of the TME at different times before and during antitumor therapy, it is possible to make prognostic statements about the effectiveness of the therapy. The described method for visualizing small pH changes in the TME of tumors differs in several aspects from previously known methods. The most used methods in this field are MR-based techniques. However, these techniques have several limitations, namely a relatively small field of view, low spatial resolution, low signal-to-noise ratio, the application of special contrast agents for *in vivo* pH imaging and expensive specialized coils and equipment (50, 59, 60).

Here studied pH sensitive [^18^F]FDG radiopharmaceuticals are easy to synthesize, are administered in the same tracer doses as [^18^F]FDG and take advantage of the good resolution of existing PET imaging technologies, which enable the imaging of large field of view.

In contrast to previously published pH sensitive radiopharmaceuticals, the uptake of [^18^F]FDG-4MBA and [^18^F]FDG-BA in the tumor is high, allowing quantitative imaging of small pH differences of the TME.

Our results show the development and application of a radiopharmaceutical consisting of a PET tracer, [^18^F]FDG routinely used in the clinic with a substituent bound to it. This radiochemical which is cleaved in a pH-dependent manner to visualize small pH differences in acidic TME. With this system, relevantly small pH differences in the acidic TME of different tumors can be visualized and due to its simple synthesis, this system might be well suited for translation into clinical trials.

## Acknowledgements

This work was supported by the Deutsche Forschungsgemeinschaft (DFG), SFB 1292 TP01 (T.Bop.), SFB 1066 project B13 (T.Bop.), B8 (T.Bop.) and B16N (T.Boh., M.M. and T.W.), TR SFB 156 B11N (T.Bop.),“Universitäres Centrum fu□r Tumorerkrankungen (UCT)”, and the ‘‘Forschungszentrum Immuntherapie (FZI)’’ of the University Medical Center.

